# Evaluating oscillatory mechanisms underlying flexible neural communication in the human brain

**DOI:** 10.1101/2025.10.16.682957

**Authors:** Varun Madan Mohan, Thomas F. Varley, Anthony M. Harris, Robin F. H. Cash, Caio Seguin, Andrew Zalesky

## Abstract

How the brain orchestrates the flow of information between its multiple functional units flexibly, quickly, and accurately, remains a fundamental question in neuroscience. Multiple theories identify neural oscillations as a likely basis for this process. However, a lack of empirical validation of proposed theories, particularly at the whole-brain scale, has hampered consensus on oscillatory principles governing neural communication, limiting our understanding of a process central to perception and cognition and its integration into experiments and clinical applications. Here, we empirically validate previously proposed neural-oscillatory communication mechanisms in the human brain – specifically those involving power and interareal phase coherence – at the whole-brain scale. We do this by estimating the dependence of inferred communication on oscillatory measures that have been theorised to facilitate communication, in source-localised resting-state magnetoencephalography (MEG) recordings. We find that power and phase coherence in the alpha, beta, and high-gamma bands track communication better than others. Crucially, the relation between communication and oscillatory measures varied across regions, indicating spatial heterogeneity in routing mechanisms. Notably, power and coherence-based principles tracked communication patterns of unimodal regions better than those of transmodal regions. In sum, these findings suggest that the human brain implements regionally specific communication mechanisms with complex neural-oscillatory dependence.

**Significance statement:** Flexible and dynamic routing of information between neural elements is vital for human brain function. Despite its importance, the mechanisms underlying flexible communication in the brain are not fully understood. Several lines of evidence suggest that neural oscillations might mediate flexible communication, but whether such mechanisms operate at whole-brain scales remains uncertain. Here, we investigate the oscillatory underpinnings of communication by assessing how well observed communication patterns from magnetoencephalography data align with the predictions of established theories. Our findings suggest that the brain follows complex region-specific routing principles to realise flexible communication. Knowledge of these principles can develop our fundamental understanding of functions like perception and cognition, refine models of interareal interactions, and help develop more effective stimulation-based clinical interventions.

## Introduction

The brain relies on efficient and effective communication, or routing of information between regions, for healthy function (Avena-Koenigsberger et al., 2018; Başar et al., 2001; Buzsáki & Draguhn, 2004; Girardi-Schappo et al., 2021; Griffa et al., 2023; Schipul et al., 2011; Schnitzler & Gross, 2005; Seguin et al., 2020; Seguin, Jedynak, et al., 2023). Neural communication is known to be flexible, or dynamically modulated by several processes, including attention, task/cognitive demands, and processing requirements (Griffa et al., 2017; Lega et al., 2016; O’Connor et al., 2002; O’Neill et al., 2015; Voloh & Womelsdorf, 2016), resulting in complex information processing pathways (Avena-Koenigsberger et al., 2019; J. R. Cohen & D’Esposito, 2016; Friederici, 2011; Hickok & Poeppel, 2007; Milner & Goodale, 1992). However, despite its fundamental functional role, the information routing mechanism that enables flexible communication in the human brain is not yet fully understood, although several experimental and theoretical advances have identified neural oscillations as a likely basis (Akam & Kullmann, 2010; Engel et al., 2001; Engel & Fries, 2010; Fries, 2005, 2015; Hillebrand et al., 2016; Odean et al., 2023; Osipova et al., 2008; Palmigiano et al., 2017; Reyes, 2003).

Neural oscillation-based theories of communication can be roughly categorised into three broad paradigms – coherence (Fries, 2005, 2009, 2015), power (Buehlmann & Deco, 2008; Jensen & Mazaheri, 2010), or resonance (Hahn et al., 2014) – based on the neural-oscillatory property purported to facilitate communication. Despite their inherent differences, these paradigms may not necessarily be incompatible with each other, as illustrated by works that attempted to unify some of them to arrive at more generalised communication principles (Bonnefond et al., 2017; Hahn et al., 2019). Several lines of experimental evidence in animal, and to a relatively lesser extent, human studies support oscillatory communication mechanisms (Bosman et al., 2012; Buehlmann & Deco, 2010; Chapeton et al., 2019; Fries, 2015; Von Stein & Sarnthein, 2000; Womelsdorf et al., 2007), although their validity at the whole-brain scale in humans has largely remained speculative. Notably, the brain is marked by clear heterogeneities in terms of function, oscillatory features, network properties, genetic makeup, directionality of information flow etc., raising the question of whether these heterogeneities extend to routing mechanisms as well. Brain regions are also regarded to have a complex functional hierarchical organisation with implications to communication (Bazinet et al., 2021; Margulies et al., 2016; Mesulam, 1998; Vázquez-Rodríguez et al., 2020). Based on this, we hypothesise that communication in the human brain might be realised through regionally specific routing mechanisms that vary across the cortical surface, and additionally across functional hierarchies, particularly along the unimodal-transmodal axis. Testing these notions would specifically require investigating previously proposed theories in the context of communication at the whole-brain scale.

In this study, we empirically validate the relationship between inter-regional communication and neural-oscillatory mechanisms at the whole-brain scale by (1) inferring communication between brain regions from neural activity time courses; and (2) correlating these to concurrently estimated neural oscillatory measures (Fig.1). We employ a previously developed method to infer inter-regional neural communication, specifically designed to track individual, directional, and high-temporal-resolution signalling events across the human brain, termed Event-marked Windowed Communication (EWC) (Madan Mohan et al., 2025). EWC infers activity propagation from neural recordings using conventional measures of functional connectivity (FC), estimated over select “communication windows” or subsamples marked by salient signal features. This approach enables it to focus on parts of the signal likely to contain instances/effects of inter-areal communication and track effects of both endogenous and exogenous perturbations. Its windowed approach also permits the simultaneous estimation of neural-oscillatory measures within the same communication windows; we specifically focus on the dependence of interareal communication on target power and source-target phase coherence (operationalised as Inter-Site Phase Clustering, ISPC).

**Figure 1.**
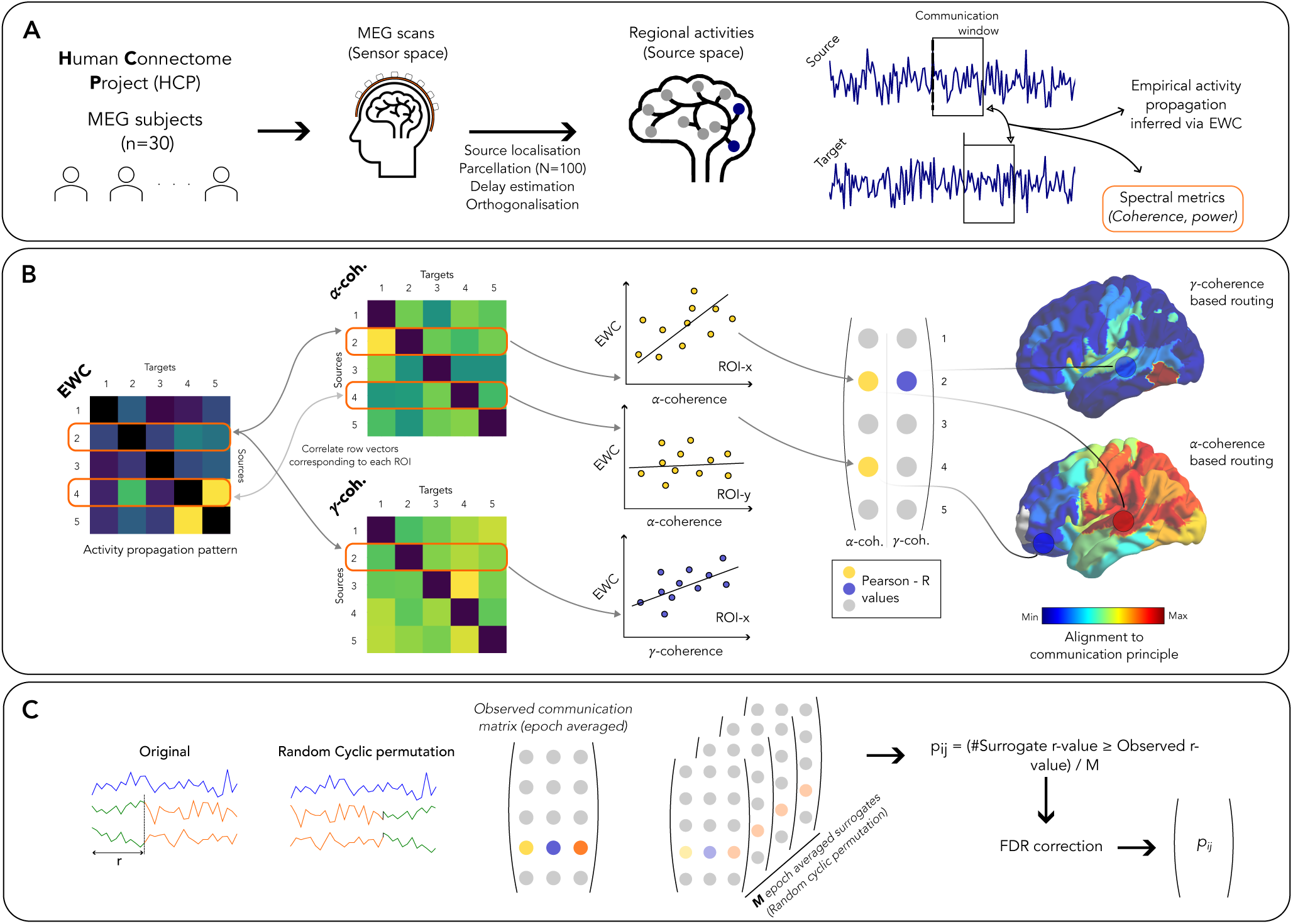
Analyses overview. (A) Preprocessing of MEG recordings – Resting state MEG recordings from the Human Connectome Project were source localised and parcellated. (B) Estimation of regional alignment to routing principles – The communication between regions was systematically inferred using Event-marked Windowed Communication (EWC). Within the same windows defined by EWC, neural oscillatory measures were estimated. The oscillatory measures were then correlated to EWC, and the correlation strengths, which captures regional alignment to a routing principle, were projected onto the cortical surface. (C) Surrogate data generation – To minimise biases originating from trivial relationships between EWC and the tested measures, a surrogate dataset was constructed aOer cyclically permuting target time series relative to the source, prior to carrying out the steps in (B).

## Results

Our primary aim in this study was to assess the validity of previously proposed neural-oscillatory mechanisms of flexible communication in the human brain, at the whole-brain scale (Fig.1). We focus on two paradigms of neural-oscillatory communication mechanisms – 1) Communication dependent on target power, and 2) Communication via phase coherence. We used source-localised resting state magnetoencephalography (MEG) recordings of 30 subjects from the Human Connectome Project (Fig.1A), from which we inferred interareal communication between pairs of brain regions (source and target) and simultaneously estimated target power and ISPC. We restrict our analyses to anatomically connected brain regions, allowing us to focus on how neural oscillations might mediate local signal transmission, which would successively build up the polysynaptic paths between mutually disconnected regions. By determining the correlation strength between communication (inferred via EWC) and power/ISPC estimates, we gauged each source brain region’s alignment to a specific communication principle (Fig.1B). These correlation coefficients were then projected onto the cortical surface for visualisation, yielding a cortical map of putative mechanisms facilitating regional communication. To ensure that the correlation coefficients did not capture a trivial dependence between inferred communication and oscillatory measures, the maps were filtered using a cyclic surrogate-based threshold (see Methods). By specifying how well the tested oscillatory measures track the information outflow from each source, these maps provide an important step towards the validation of mechanisms of flexible information transfer in the human brain.

### Dependence of communication on target power

We first set out to characterise the relationship of EWC between pairs of regions (a source and target) to the target’s oscillatory power in the theta, alpha, beta, and gamma (-low and - high) bands (Fig.2A).

**Figure 2.**
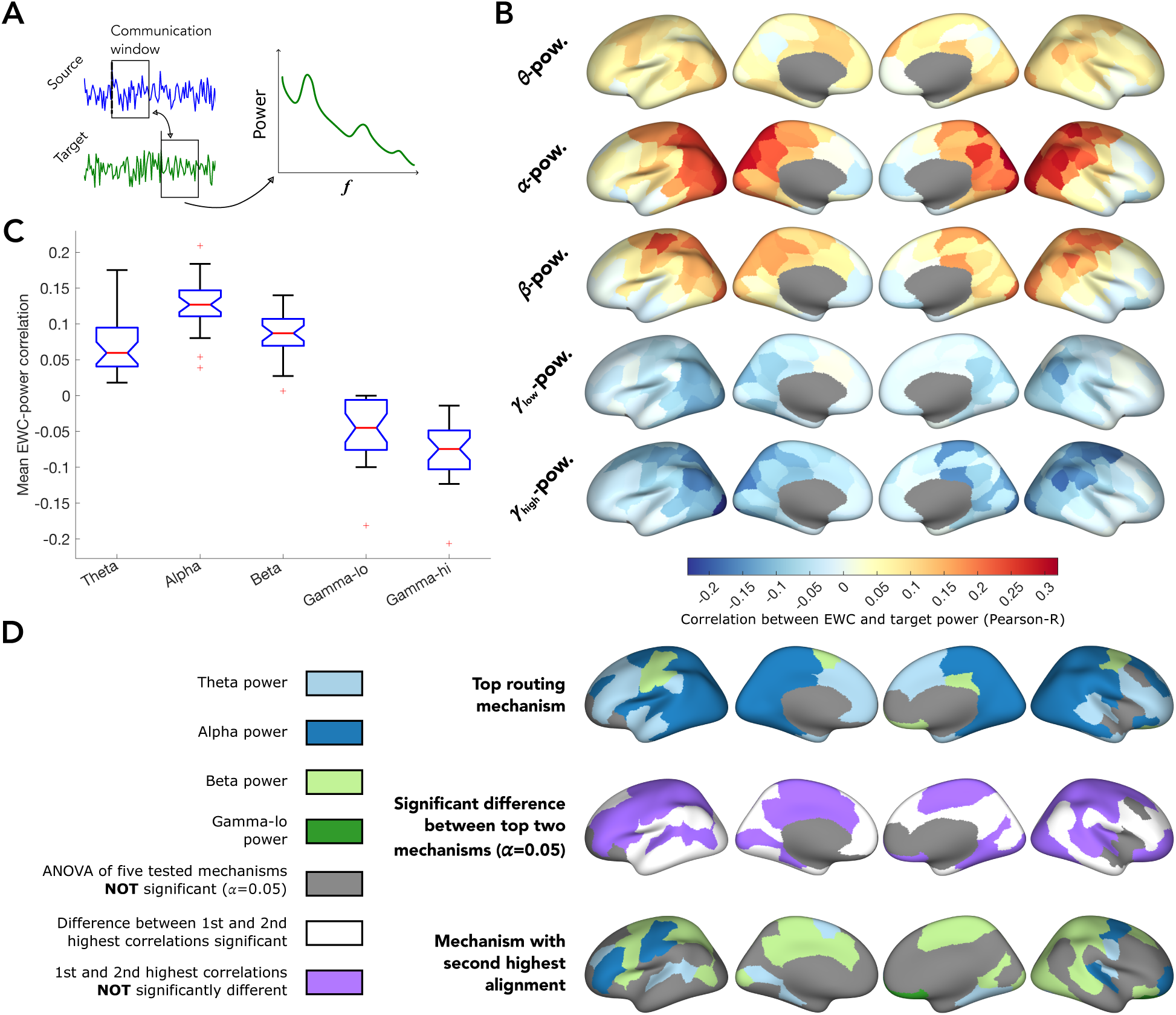
Dependence of communication on target power. (A) For a source and target pair, along with the EWC, the target’s power spectral density is estimated for the signal contained in the communication window. (B) The correlation between EWC and target power in each frequency band is projected onto the cortical surface. The colour captures how strongly information flowing out of a region is correlated to target power. (C) Distribution (across all regions) of correlation strengths, to target power in each frequency band. (D) (top) Frequency band in which target power maximally correlates with EWC; (middle) Binary classification based on whether the top two correlations are significantly different (α=0.05). (bottom) Frequency band of the second highest correlation, displayed only in regions where the difference between the top two bands is not significant.

We observed a marked spatial heterogeneity in EWC’s dependence with target power in each of the frequency bands, indicative of a location-specific routing mechanism (Fig.2B). Specifically, the dependence of outward information flow on both target alpha and beta power was maximal in the posterior and parietal brain regions typically associated with visual and somatomotor areas, and predominantly positive (Fig.2C). Interestingly, the dependence of inferred communication on both gamma-lo and gamma-hi was also maximal in the posterior regions, but in the negative direction (Fig.2C), indicating that information flow out of those regions were directed to neighbours with low gamma power.

The distribution of the top oscillatory measure (Fig.2D-top) reveals a clear variation of the routing principles along the anterior-posterior axis, with information flow from anterior regions depending maximally on the target theta-power, and communication originating from posterior regions depending maximally on alpha-power, and a few regions exhibiting beta power dependence. We then tested whether there was a significant difference between the correlations obtained for the two top mechanisms of each region (Materials and Methods). This step was important to identify regions for which a single oscillatory measure stood out in explaining neural communication inferred using EWC. We found that the correlation of the top routing principle was statistically similar to the second-best routing principle (beta power in most cases – Fig.2D-centre; regions in red) in a large number of regions. However, there were also several regions – located predominantly in medial prefrontal, temporal and occipital areas – whose communication patterns were best explained by target power in a single frequency band (Fig.2D-centre; regions in white). Notably, we found only one region, the right orbital frontal cortex, where gamma band power, specifically gamma-lo, predicted routing.

In short, our investigation into information routing in the brain based on power reveals a rich heterogeneous landscape of the relationship between communication and target power. We find that the dependence of communication on power varies not only with the frequency, but also spatially, across the cortical surface. This indicates that signalling originating from different regions might be facilitated by different oscillatory mechanisms. We also find evidence suggesting that a routing principle based on oscillations in a single frequency band might not fully capture empirical communication mechanisms.

### Dependence of communication on phase coherence

In our second set of analyses, we investigated another paradigm of neural-oscillatory mechanisms – communication through coherence. Like the approach in the previous analyses, here we correlated inferred communication patterns captured by EWC, to the Intersite Phase Clustering (ISPC) between sources and anatomically connected targets in the five frequency bands, within the same communication windows (Fig.3A): ISPC captures the consistency between phases of a pair of signals, indicating the level of synchronization between them (see Methods). Projecting the correlation values onto the cortical surface, we found that the dependence of communication on ISPC followed a spatially heterogeneous profile as in the case with target power (Fig.3B). For instance, while both alpha and beta ISPC showed similar correlation to EWC across regions (Fig.3C), the regions associated with alpha coherence were primarily localised in the posterior part of the brain, whereas beta coherence best predicted EWC in parietal and somatomotor regions. Across regions, ISPC in the alpha, beta, and gamma-hi bands correlated with EWC better than theta and gamma-lo bands (Fig.3C).

**Figure 3.**
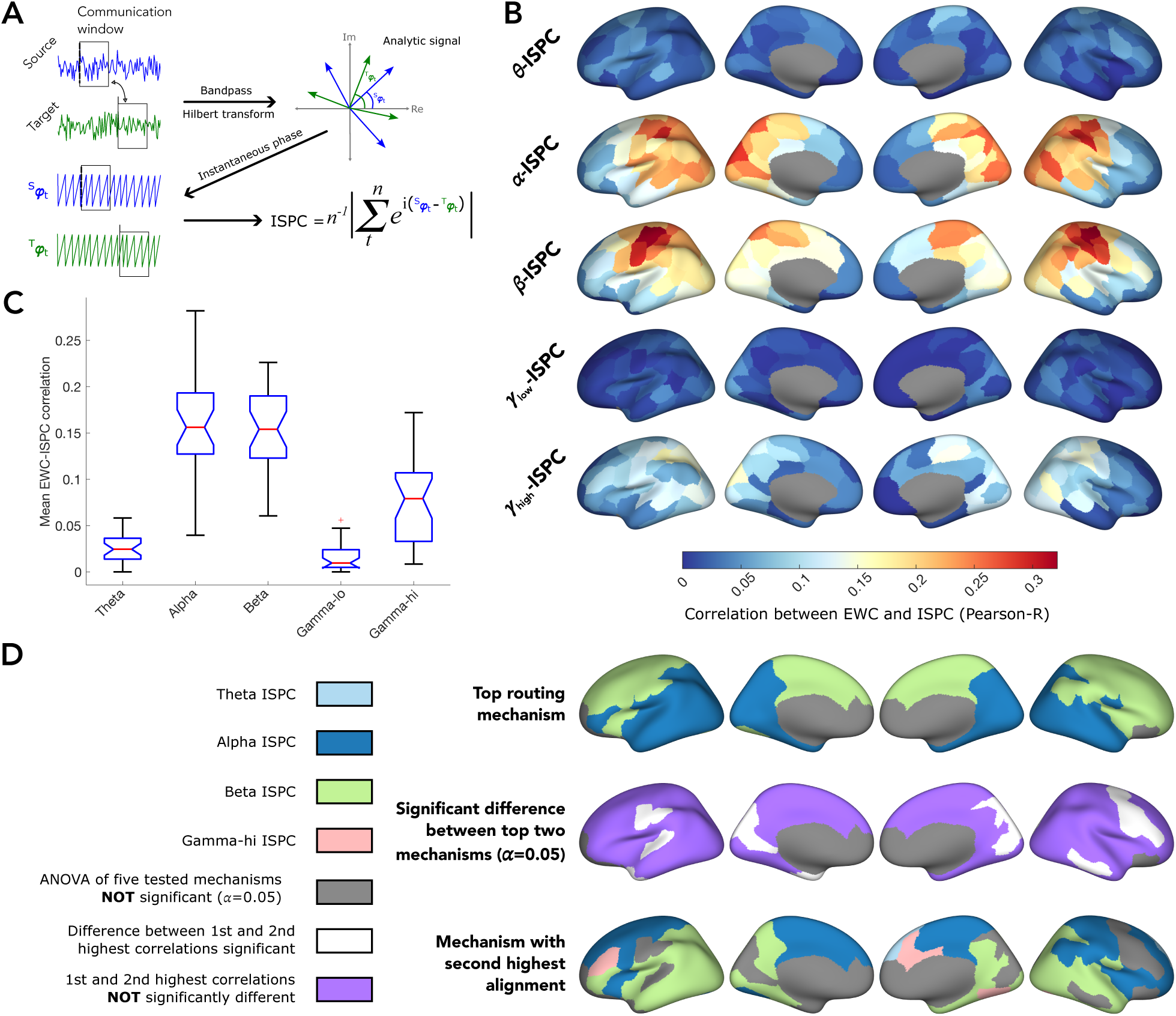
Dependence of communication on coherence. (A) For a source and target pair, the signals are first bandpass filtered into the theta, alpha, beta, gamma-lo and gamma-hi bands. The phase time series is then extracted from the signals aOer Hilbert transform. The Intersite Phase Clustering (ISPC) is then estimated from the phase time series, along with EWC, which is estimated from the raw signals. (B) The correlation between EWC and ISPC in each frequency band is projected onto the cortical surface. The colour captures how strongly information flowing out of a region is correlated to ISPC. (C) Distribution (across all regions) of correlation strengths, to ISPC in each frequency band. (D) (top) Frequency band in which ISPC maximally correlates with EWC; (middle) Binary classification based on whether the top two correlations are significantly different (α=0.05). (bottom) Frequency band of the second highest correlation, displayed in only those regions where the difference between the top two bands is not significant.

As in our analyses with target power, for each region, we identified the top two bands in which ISPC maximally correlated with its EWC and tested whether the correlations were significantly different. As argued previously, this would help ascertain whether only a single oscillatory measure could predict a region’s communication with its neighbours. Although looking at measures with the maximal correlation to EWC clearly revealed alpha ISPC dominance in posterior inferior regions and beta ISPC dominance in the anterior regions (Fig.3D-top), we found that in most regions, the second highest correlation was statistically similar to the first (Fig.3D-middle), suggesting that ISPC in at least two bands similarly tracked EWC patterns. Regions in which ISPC in a single band tracked EWC better than others were much fewer, located in the medial occipital, left superior temporal and central, and right inferior temporal and frontal areas. It was further interesting to observe that in most regions where the maximal EWC correlation was to alpha ISPC, the second highest correlation was to beta ISPC, and vice versa (Fig.3D-bottom).

To summarize, in this set of analyses, we investigated the degree to which whole-brain communication aligned with coherence-based information routing principles. We found that communication, as captured by EWC, has a notably heterogeneous dependence on coherence in the different bands along the cortical surface. Additionally, the cortical distribution also varies with the frequency band, indicating regional specificity in the routing principle followed. ISPC in the alpha and beta bands correlated the most strongly with EWC. We further found that in most brain regions, ISPC in at least two frequency bands could similarly predict information flow captured by EWC.

To ensure that the heterogenous dependence of EWC on power and coherence was not specific to our choice of the dataset, we carried out similar analyses on a separate dataset, with different acquisitional parameters and parcellation (Fig.S2). We note that the regional dependence of EWC on both ISPC and power in all bands follow a qualitatively similar spatial profile as in Fig.3.

### Neural oscillatory dependence of communication across functional gradients

Our previous analyses revealed clear spatial and spectral heterogeneity of inferred communication’s dependence on power and ISPC. The parietal and posterior localisation of the regions showing maximal correlation to the oscillatory measures, however, prompted us to explore whether the extent to which these measures tracked communication varied with gradients in functional organisation, particularly about the unimodal-transmodal axis (Margulies et al., 2016; Mesulam, 1998). In our final set of analyses, we estimated the degree of spatial alignment between the EWC-ISPC correlation maps obtained in our previous analyses and the principal functional gradient derived from group-level FC maps (Fig.4).

**Figure 4.**
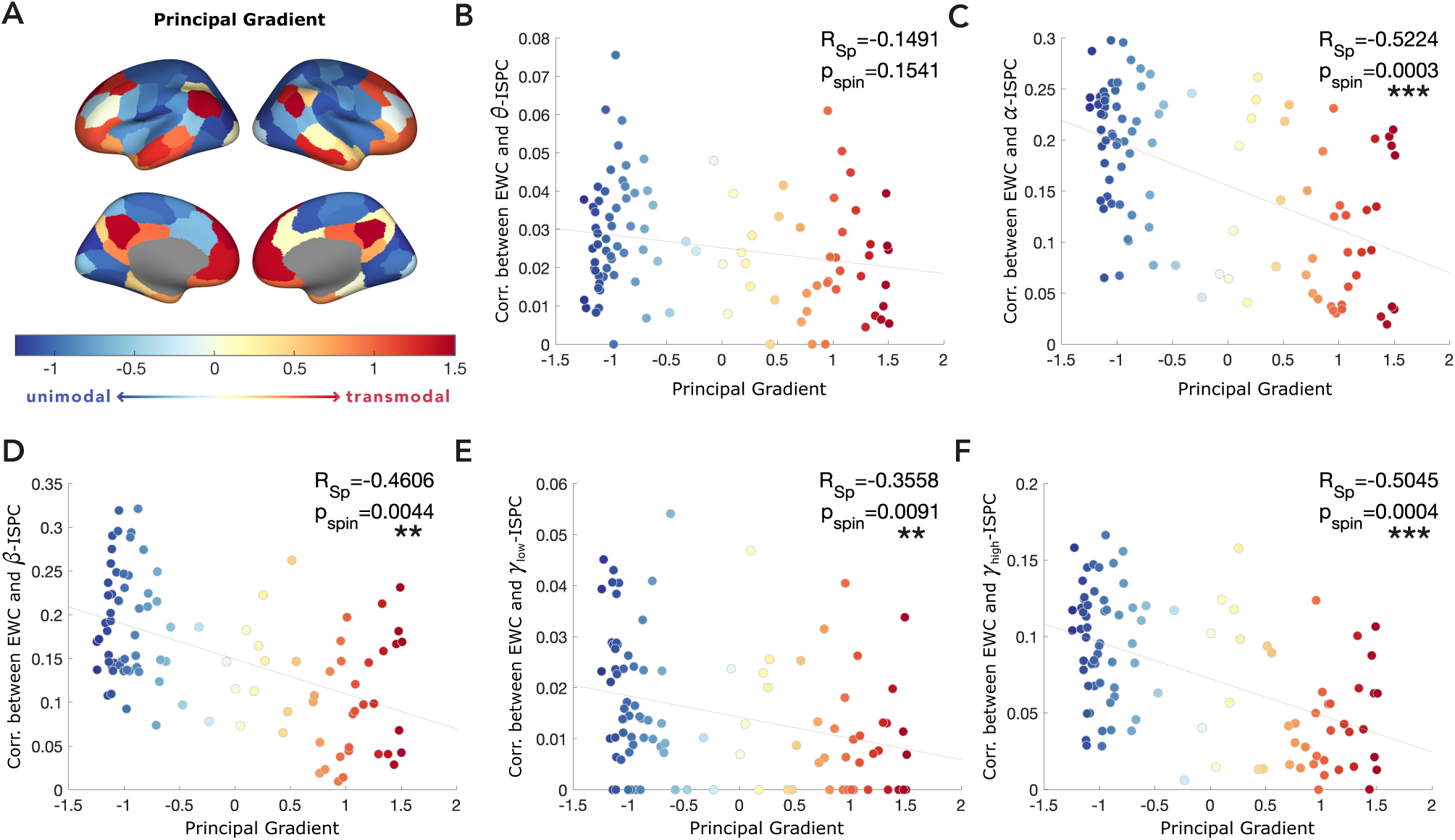
Strength of the correlation between a EWC and ISPC depends on the source region’s placement along the principal functional gradient. **(A)** Principal functional gradient derived from diffusion embedding of group-averaged FC. **(B-F)** Correlation between EWC and ISPC in the frequency bands of interest, as a function of the source region’s principal gradient coefficient (also represented in the colour of the point). Each point represents a brain region. Relationship to functional gradient is quantified as the Spearman correlation, and p-value is obtained by comparison to surrogate dataset with 10000 spin permutations.

We also estimated the gradient’s alignment to the EWC-power maps (Fig.S3). We observed that the correlation strength between inferred communication and coherence varied across the principal functional gradient delineating the unimodal-transmodal axis, for the alpha (Spearman R, *R_sp_* = −0.5224, spin permutation-based *p_spin_* = 0.0003), beta (*R_sp_* = −0.4606, *p_spin_* = 0.0044), gamma-lo (*R_sp_* = −0.3558, *p_spin_* = 0.0091) and gamma-hi (*R_sp_* = −0.5045, *p_spin_* = 0.0004) bands. Interestingly, the variation across the gradients for all bands revealed that the correlation between ISPC and communication was stronger for unimodal regions compared to transmodal regions. This difference was most prominent for the alpha band, whereas the dependence of communication on theta ISPC did not show significant variation across the unimodal-transmodal axis. Like ISPC, power also tracks communication primarily in unimodal regions Fig.S*3*.

To summarise, in this section we examined the utility of ISPC and power in explaining a region’s communication patterns given its position in the functional unimodal-transmodal axis. We found evidence suggesting that communication based on both phase coherence as well as power, particularly in the alpha, beta, and high gamma bands is more dominant in unimodal regions rather than transmodal regions.

## Discussion

The mechanisms and principles underlying communication over different scales, that enable the brain to perceive, process and act on stimuli within milliseconds, have understandably been a topic of much interest (Avena-Koenigsberger et al., 2018; Lerner et al., 2016; Roland et al., 2014; Seguin, Sporns, et al., 2023). Over the years, there has been an accumulation of a wealth of experimental data, insights and theories of neural communication, and the development and refinement of biophysically accurate models, allowing us to develop means to observe and study brain communication in great detail. Although neural oscillations are widely considered to underpin mechanisms of flexible communication, there is a lack of consensus on the exact mechanism that operates in the human brain and whether the oscillatory dependence is consistent across regions. In this work, we set out to address this outstanding question and concurrently place established communication theories in the more general context of information transfer at the whole-brain scale.

We specifically explore two flexible oscillatory communication paradigms – communication dependent on target power, and communication through phase coherence. To do this, we first inferred the communication between anatomically connected regions and simultaneously estimated oscillatory measures associated with proposed theories. This was followed by estimating the correlation between them, effectively capturing the alignment of regional communication towards putative routing mechanisms. Across both the tested paradigms, alpha and beta band oscillations were the most associated with communication (Fig.2C,3C). In the case of EWC-power correlations, it could be argued that the observations result from inherent power heterogeneities and non-uniform spectral densities. However, the same cannot be said for EWC-ISPC correlations since the ISPC is estimated on the instantaneous phase time series of a band-passed signal and effectively power normalised (Bruña et al., 2018; M. X. Cohen, 2014). Despite this, it was noteworthy that the observed heterogeneities in ISPC-dependence seemed to match inherent regional power heterogeneities, particularly in the alpha (posterior dominant) and beta (dominant in parietal somatosensory areas) bands. This suggests that regional power distributions may play a role in determining what oscillation-based routing mechanism is followed by a region. Additionally, it is interesting to observe that in both the paradigms, in most regions, the top two oscillatory measures track EWC in a statistically similar manner (Fig.2D,3D). This suggests that empirical communication mechanisms might involve multiple frequency bands, as opposed to communication over a single band. A routing mechanism based on the observed heterogeneous oscillatory dependences would imply that regions locally communicate via the same frequency band, but over different frequency bands for distant targets, in a manner similar to the distance-based routing mechanism earlier proposed by (Von Stein & Sarnthein, 2000).

In our analyses of the dependence of inferred communication (via EWC) on target power, it was interesting to observe the positive association of EWC to target alpha power and the negative association to target gamma power. At first glance these observations seemed to be in contradiction to literature linking alpha to functional inhibition and gamma activity to neural processing which would suggest that the associations be sign-flipped (Jensen & Mazaheri, 2010). A better interpretation of our observations requires a closer look into EWC and what it measures: the lagged partial correlation between source and target signals, essentially capturing the degree of synchronisation between the signals in the time domain.

From a signal processing standpoint, an increase in the high frequency gamma band component of the target would imply stronger rapid fluctuations of the signal and might serve to reduce synchronisation, effectively resulting in a negative association. On the other hand, an increase in inhibition, which is well associated to increases in alpha power, is also counterintuitively linked with an increase in synchronisation (Hanslmayr et al., 2012; Klimesch et al., 2007; Van et al., 1994) which might explain the positive association between EWC and target alpha power. Future work combining neurotransmitter imaging with localised neural recordings, or using a measure for differential neural processing capabilities between pairs of regions as a proxy for communication will be able to establish this link more definitively. Our seemingly counter-intuitive observations also help remind us that the low-level signal features we observe do not always map directly to cognitive/behavioural implications, which often require several additional levels of insights and inference.

Our results also revealed that communication in the anterior regions, often associated with higher order brain function, showed negligible oscillatory dependence in the case of power, and only a weakly positive dependence with ISPC in the alpha, beta and gamma-hi bands (Fig.2B,3B). This was further emphasised in the functional gradient analysis, where the coherence and power dependence in transmodal regions were found to be lower than in unimodal regions (Fig.4). This could suggest that higher-order regions might follow a more complex oscillatory, or perhaps even non-oscillatory routing mechanism, to facilitate the integration of information from various input streams.

Testing the oscillatory dependence of communication in an empirical setting is advantageous on many fronts. For instance, knowledge of the principles describing communication dynamics will help us better understand how the underlying structural connectivity (SC) relates to observed FC (Avena-Koenigsberger et al., 2018). On the computational modelling front, the incorporation of flexible communication will make models of whole-brain activity more realistic. It can also help build intuition into communication dynamics associated with neural oscillatory changes linked to cognitive task processing and neuropathologies. In a clinical setting, understanding the principles of communication dynamics can help in predicting and effectively controlling information flow through stimulation-based interventions such as transcranial magnetic stimulation (TMS) or deep brain stimulation (DBS). By capturing the regional variations in the alignment of communication to different frequency bands, our results provide empirical support for existing communication theories involving various oscillatory measures and frequencies (Bonnefond et al., 2017; Buehlmann & Deco, 2010; Chapeton et al., 2019; Engel & Fries, 2010; Fries, 2005, 2015; Jensen & Mazaheri, 2010; Palmigiano et al., 2017) in a whole-brain context, while also identifying potential limitations to their validity. A future direction of research could explore how we can combine all the correlation strength maps to arrive at a comprehensive communication principle for the cortical surface. One possible model would be a winner-takes-all approach, where the communication principle of a region is defined as the spectral metric that correlates the most with communication. The most important assumption there would be that the other spectral metrics might not facilitate communication, or have negligible contributions compared to the metric that is the most correlated with communication strengths. A model that would not discount the contributions of the other bands might be one in which the routing principle is the sum of the various spectral metrics weighted in proportion to their location specific correlation strengths. Furthermore, although we only explore the dependence of communication on power and phase coherence in this work, future work could also test the validity of other oscillation-based theories such as communication through resonance (Hahn et al., 2014), or those simultaneously involving multiple frequency bands and measures (Bonnefond et al., 2017; Hahn et al., 2019).

Finally, an alternative approach to conceptualising how information is routed between regions is through network-based models, where the white matter, or structural connectivity between regions, determines the route and efficacy of information transmission between them. These models have been shown to adequately predict function and even stimulus propagation paths (Seguin et al., 2020; Seguin, Jedynak, et al., 2023; Seguin, Sporns, et al., 2023). Flexible information flow can also be realised through a dynamic anatomical substrate (Pope et al., 2023). The network-based and oscillation-based paradigms are not mutually exclusive, but rather capture different aspects of the routing process. While network-based models describe possible communication routes between regions for a given network, oscillation-based models describe the mechanism by which those routes may be realised. Bridging these paradigms, by exploring oscillation-based routing principles as a generative model for network-derived communication routes, could be an important direction for future research. *Limitations* While our results suggest that information transfer in the human brain follows heterogenous routing principles, it is essential to consider the analytical and methodological limitations of this study to better contextualise our findings. Firstly, it is important to mention that our notion of information transfer / communication is statistically inferred from neural activity time courses – it is not a direct measure of regional responses to a perturbation, as in the case of localised neurostimulation-response measurements. Secondly, the need for non-invasive whole-brain sub-second resolution neural dynamics drove our choice for source-localised MEG to study the principles of endogenous communication. Although source localisation yields ROI-level recordings, the reduced spatial resolution of MEG can cause sensor-level signals to be “smeared” across sources, and subject to assumptions of the source localisation technique used. If the spatial extent of the investigation is reduced, more spatially localised modalities such as intracranial EEG (iEEG) would help bypass several such limitations. Finally, it is crucial to acknowledge that although we had to infer communication as well as estimate the neural oscillatory measures from the same underlying signal for the current study, requiring us to employ a surrogate-based correction of observed relationships, an ideal approach would be to estimate these measures separately from concurrent and independent modalities when available.

## Conclusion

In conclusion, our work explores the validity of neural oscillation-based mechanisms of communication at the whole-brain scale. We find that measures like power and phase coherence, theorised to facilitate communication processes, track communication in a regionally specific and hierarchical manner. Our findings shed light on the complex neural oscillatory underpinnings of communication in the brain.

## Methods

### Dataset

#### Human Connectome Project (HCP)

Resting state MEG scans of 30 subjects (22-35 years, 17F), along with associated MEG anatomical data, 3T structural MRI data, and empty-room recordings, were obtained from the Human Connectome Project (Van Essen et al., 2013), through the ConnectomeDB platform. MEG was acquired using a 4D Neuroimaging MAGNES 3600 system at a sampling rate (SR) of 2035Hz (Larson-Prior et al., 2013). Recordings varied in duration from 5-6 minutes, and anti-aliasing low pass filtered at 400Hz.

#### Open MEG Archive (OMEGA)

To test the generality of our findings beyond one dataset and set of acquisition parameters, we carry out all our analyses on a second dataset - a subset of the Open MEG Archive (OMEGA) dataset. Resting state MEG scans of 5 subjects (21-35 years, 2F), along with 3T structural MRI data, and empty-room recordings collected as a part of the OMEGA initiative, was obtained from OpenNeuro (dataset version 1.0.2 : https://dx.doi.org/10.18112/openneuro.ds000247.v1.0.2) (Niso et al., 2016). The recordings were acquired using a CTF 275 system with a SR of 2400Hz. Recordings were 5 minutes long, and anti-aliasing low pass filtered at 600Hz.

### Processing

The MEG recordings from both HCP and OMEGA datasets were processed entirely using the Brainstorm software (Tadel et al., 2011) on MATLAB (The MathWorks Inc., 2022), largely in accordance with the pipeline described in Brainstorm tutorials (Niso et al., 2019). In the HCP data, MEG recordings were first coregistered to the subject’s structural MRI using the MEG anatomical data. We did not have access to the fiducial positions for the OMEGA dataset and thus defined approximate fiducial locations which were then refined using the digitised head points. A notch filter (60, 120, 180, 240 and 300Hz), followed by high pass filter (0.3Hz) were applied to resting state and empty-room recordings, to filter out power-supply and slow-wave/DC-offset artifacts respectively. Each subject’s recording was then visually inspected, along with the channel power spectral density, to weed out bad channels and bad time segments. The ECG and EOG recordings were then used to identify heartbeats and eye-blinks, after which the identified artifacts were removed using their signal space projections (SSPs) (Uusitalo & Ilmoniemi, 1997).

Source-level activities were then estimated from sensor-level recordings at 8004 points corresponding to the fsLR4k mesh for the HCP data, and 15002 points for the OMEGA data. This involved first computing the head model using overlapping spheres and constrained dipoles normal to the cortical surface, and estimating the noise covariance from the empty-room recordings. Source-level activities were then estimated using the dSPM method (Dale et al., 2000) available in Brainstorm. The sources were then parcellated using the Schaefer-Yeo 7-network 100 atlas (N=100) (Schaefer et al., 2018) for HCP and Destrieux atlas (N=150) (Destrieux et al., 2010) for OMEGA, with the parcel activity computed as the principal component of the constituent source activities.

#### Structural connectivity

Structural brain networks for 1000 healthy young adults from the HCP were mapped from minimally preprocessed high-resolution diffusion-weighted MRI (Glasser et al., 2013). Following previous work (Mansour L et al., 2021; Seguin et al., 2022), an MRtrix3 probabilistic tractography pipeline (Tournier et al., 2012) was used to map whole-brain white matter tractograms - multi-shell multi-tissue constrained spherical deconvolution (Tournier et al., 2007) iFOD2 tracking algorithm (Tournier et al., 2010), anatomically constrained tractography (R. E. Smith et al., 2012), 5M streamlines - see (Mansour L et al., 2021) for further details. Cortical gray matter was parcellated using the Schaefer-Yeo 7-network 100 atlas (Schaefer et al., 2018). Connection weight between pairs of regions was defined as the number of streamlines normalised by the product of their surface areas. Connections comprising less than 5 streamlines were discarded to minimise the false positive rate of probabilistic tractography (Zalesky et al., 2016). A distance-dependent consensus algorithm was used to combine subject-level structural networks to a group-level structural connectome (SC) (Betzel et al., 2018). A proportional threshold was then applied to the group-level SC to retain only the top 15% structural connections, the SC was binarised, and inter-hemispheric connections were removed. This SC was used for the analysis of the MEG recordings from the HCP dataset (source localised and parcellated using the same atlas), to identify the anatomically connected neighbourhood of each source.

DWI-derived subject-level SCs of 50 healthy young adults in the Destrieux parcellation (N=150) were obtained from the publicly available MICA-MICs dataset (Royer et al., 2022). As done previously, we combined the subject-level SCs using a distance-dependent consensus algorithm to obtain a group-level SC, which was proportionally pruned to retain the top 15% connections. The SC was then binarised, inter-hemispheric connections were removed, and used in the analyses of the OMEGA dataset’s MEG recordings (source localised and parcellated using the Destrieux atlas).

#### Functional connectivity

We obtained minimally preprocessed ICA-FIX resting-state functional MRI data of the same 1000 healthy young adults from the HCP. Four scans (2 sessions: right-to-left and left-to-right phase encoding scans, on 2 separate days) spanning 14 minutes and 33s (TR=0.72s) were collected for each participant, for a total of 4000 subject-level scans (Acquisition protocols detailed in (Glasser et al., 2013; S. M. Smith et al., 2013)). The time series of voxels belonging to the same gray matter region delineated by the Schaefer-Yeo 7 Network 100 atlas were averaged, and the functional connectivity (FC), corresponding to each scan, was computed as Pearson correlation between all pairs of regional time series. Group-level FC was computed as the average of the 4000 subject-level FCs (Seguin et al., 2022).

### Inferring inter-areal communication

We used a previously developed method to infer communication between regions, termed Event-marked windowed communication (EWC). EWC uses a targeted, temporally ordered, windowed approach to gauge the transmission of endogenous perturbations between neural elements from their activity time series (Madan Mohan et al., 2025). Briefly, EWC involves (1) the identification of salient features of the source region’s signal, which are termed “events”; (2) the definition of a subsample of the signal at the location of the events, and at temporally displaced locations in the target’s signal, proportional to the conduction delay between source and target; and (3) the estimation of the statistical dependence or functional connectivity (FC) between the source and target using measures such as Partial correlation, conditional mutual information etc. Thus, EWC effectively infers the propagation of endogenous perturbations by tracking lagged statistical dependencies between source-target pairs.

Prior to EWC estimation, recordings were epoched into smaller 10s segments to enable efficient computation, and each epoch was processed independently. For each epoch, event identification was done using a z-score based threshold (Sorrentino et al., 2021) of |*z*| > 3.

We follow the approach and operational parameters in (Madan Mohan et al., 2025), but summarise the steps below. The following steps were carried for each event:

1. A region was chosen as a source, and a communication window of 1*s* was defined starting at the event. After identifying the regions that were anatomically connected to the source based on the structural connectivity matrix (neighbours), a window of similar length was placed at their time series, at delayed timepoints (delay proportional to the Euclidean distance between the source and the target).
2. Partial Correlation was used to gauge statistical dependence between the source and target recordings contained in the windows. The past activity of regions (up to window length – 1s) was used as the conditional variable, to discount the effects of past internal dynamics on inferred communication. When used with partial correlation, EWC has also been shown to be highly correlated with the Transfer Entropy (TE) and bivariate Granger Causality (GC) (Madan Mohan et al., 2025).
3. The partial correlation estimates were checked for significance, with a Bonferroni-corrected threshold of 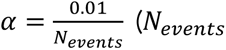 is the total number of events within the epoch), since the correlation is tested multiple times within an epoch. Estimates were set to zero above this threshold.

The EWC protocol resulted in a *N* × *N* × *M* communication matrix for a recording split into M epochs, for each subject.

Temporal ordering was proportional to the physical Euclidean separation between regions defined using centroid coordinates of the Regions of Interest (ROIs) in the Schaefer 7-network 100 atlas (for HCP) and the Destrieux atlas (for OMEGA) (Abeysuriya et al., 2018; Deco et al., 2009; Papadopoulos et al., 2020). Inter-regional distances were proportionally converted to time delays (*δ*) as

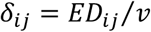

Where *v* is the conduction velocity (CV) of neural signals, set to 10 *m*⁄*s* (Papadopoulos et al., 2020). Parcellated source-localised resting-state MEG recordings were corrected for source leakage effects by removing zero-lag correlations as per (Colclough et al., 2015), using the OHBA Software Library (OSL) package in Python (Quinn et al., 2023).

### Neural-oscillatory measures

For each source-target pair of regions, the target power and phase coherence were computed alongside the EWC, for each pair of communication windows situated at the events i.e. each EWC estimate, which captured the information flow from a source to a target over a certain window of time, had an associated value of target power or phase coherence (Fig.1A).

#### Target power

To test whether the spectral power of a target influenced how much information it received from a source, for each source-target pair, we first measured the power spectral density (PSD) of the target signal contained within the communication window associated with each event. This was done using Welch’s periodogram method in MATLAB (function: pwelch), with a window size of *SR*/2. The power in each frequency band – theta (4-6Hz), alpha (8-12Hz), beta (14-24Hz), gamma-lo (30-59Hz) and gamma-hi (60-80Hz) was estimated by first integrating the PSD over the constituent frequencies through trapezoidal numerical integration, and then normalising the estimate by the total power of the PSD to give the relative power of each band.

Similar to the output of the EWC protocol described above, for each source, we obtained a value of N target powers, resulting in a final *N* × *N* × *M* matrix per subject.

#### Intersite Phase Clustering (ISPC)

To test whether the information transfer between a source and target was dependent upon the phase coherence between them, we measured the intersite phase clustering (ISPC) between the instantaneous phase time series of band-passed source and target signals contained within communication windows. ISPC, or simply the phase coherence or phase locking value, is a measure of consistency of the phase relationship between signals (M. X. Cohen, 2014; Lachaux et al., 1999), varying from 0 to 1, indicating random/independent phase relationship to perfectly consistent phase difference over time respectively. Between regions with high ISPC, the communication through coherence hypothesis (Fries, 2005) posits that information transfer should be highest for pairs regions with delay-corrected angular differences close to 0, due to their temporally coordinated excitability cycles, and lowest for those that are consistently out of phase (angular differences close to pi).

Operating on the instantaneous phase time series, ISPC computation requires Hilbert transformation of the band-passed raw signal to create a complex-valued analytic signal. Given a pair of phase time series *ϕ_x_* and *ϕ_y_*. in a frequency band f, ISPC is computed as:

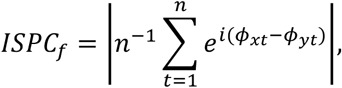

where *n* is the number of time points. ISPC can alternatively be viewed as an amplitude normalised version of the spectral coherence obtained from the Fourier transform of the cross spectral density (Bruña et al., 2018). Like EWC and power, ISPC computation between all source-target pairs resulted in an *N* × *N* × *M* matrix for each subject.

### Communication principles

To gauge the dependence of inferred inter-areal communication on the power and phase coherence, for each subject, the Pearson correlation between each row of the EWC matrix concatenated over epochs (resulting in a 1 × *NM* vector which corresponds to a single source) and the matching epoch-concatenated row in the power and ISPC matrices was estimated. This resulted in 10 Pearson coefficients per region (5 bands of power and 5 bands of ISPC) indicating how aligned the communication from that region (source) was to a certain communication principle. Estimating the correlation of all regions similarly yielded a *N* × 10 matrix per subject, which we term the observed “principle matrix” (Fig.1B).

#### Surrogate data analysis

When investigating the validity of a communication mechanism by assessing the relationship between communication and the measure theorised to underpin it, it is vital to ascertain whether the measures might have some trivial or “default” dependence between them. This is particularly important in this work, since both the measure of communication and power/ISPC are derived from the same underlying MEG signal (Fig.S1). To establish whether the observed relationship between EWC and power/ISPC significantly exceeds the trivial dependence between the measures, the elements of the *N* × 10 principle matrix capturing a subject’s regional alignment of communication to the tested spectral measures (observed correlation between EWC and power/ISPC) was compared to a cyclically permuted surrogate distribution (Fig.1C). The surrogate distribution was generated and compared to the observed principle matrix as follows:

1. We generated a surrogate dataset using a random cyclic permutation approach to shuffle the parcel time series: (A) in each epoch, a random integer is chosen between 0 and the total length of the data contained in the epoch. (B) For each source region, the activities of all other regions (targets) are cyclically permuted in the forward direction by the number of steps given by the random integer. (C) The steps in the above sections are carried out on the permuted data. By cyclically permuting the signals relative to the source, we effectively compute the EWC and neural oscillatory measures between random pairs of windows – any trivial dependence between EWC and the target power or ISPC would persist between these random windows and manifest as the (surrogate) correlation between EWC and power/ISPC.
2. An *N* × 10 surrogate principle matrix is obtained for each surrogate (the ROI-level correlation between EWC and each of the neural oscillatory measures).
3. The communication protocol is carried out for *S* (=1000) separate surrogate datasets, resulting in a total of *S* surrogate principle matrices (each with dimensions *N* × 10) for each subject.
4. The p-value for principle *l* of region *i* was defined as:

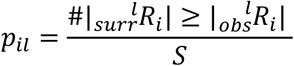 Where 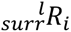 and 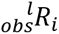 represent region *i*’s surrogate and observed Pearson correlation coefficients respectively for principle *l*, and #(…) denotes the number of instances the argument is true. If the observed dependence between EWC and the neural oscillatory measures simply reflects the trivial dependence between the measures, the p-value will not pass the significance threshold.
5. The p-values were False Discovery Rate (FDR) corrected across rows using the Benjamini-Hochberg linear step-up procedure implemented in MATLAB (function ‘mafdr’).
6. A subject-level mask with dimensions *N* × 10 was defined, such that *mask_il_* = 1 if *p_il_* < 0.05. This mask was applied to the observed *N* × 10 principle matrix of the subject.

The surrogate-corrected *N* × 10 subject-level principle matrices were averaged across subjects and projected to the cortical surface (Fig.1B).

### Functional gradient analysis

Functional gradients were computed through dimensionality reduction of group-level FC (derived from resting state functional magnetic resonance imaging) from the HCP, using diffusion embedding. We use the principal gradient or eigenvector of the diffusion operator, which varies across the unimodal to transmodal axis. The sign of the obtained gradients were flipped to enable comparison to existing works (Margulies et al., 2016). This method was implemented using the Dimensionality Reduction Toolbox (Van Der Maaten et al., 2009).

To establish whether alignment to the tested measures varied based on where a region was situated on the unimodal-transmodal gradient, we estimated the Spearman correlation strength between the principal gradient values and the EWC-power/ISPC alignment (elements of the surrogate corrected principle matrix).

### Statistical analyses

Correlations between EWC and neural oscillatory measures for each region were tested for statistical significance using a cyclic-surrogate-based permutation approach (see ‘Surrogate data analysis’). For each subject (*N* = 30 for the HCP dataset and *N* = 5 for the OMEGA dataset), we created a surrogate dataset with *S* = 1000 surrogates. Each cyclic surrogate involved cyclically shifting the MEG recording data of targets by a random amount with respect to each source region’s recording, after which correlations between EWC and neural oscillatory measures were estimated. A p-value was defined by comparing each surrogate correlation strength with the observed as follows: 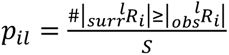, for a region *i*, and neural oscillatory principle *l*. To account for multiple comparisons, the p-values were then FDR corrected using the Benjamini-Hochberg Linear step-up procedure across regions (*N_regions_* = 100 for HCP and = 150 for OMEGA). The correlation values were then thresholded based on the FDR-corrected p-values (*⍺* = 0.05), and set to 0 if non-significant (indicating no relationship between EWC and the oscillatory measures).

To identify the top oscillatory principle for each region, we picked out the oscillatory measure which maximally correlated with each ROI’s information outflow (Fig.2D-top, 3D-top). To ensure that the top oscillatory principle explained communication significantly better than the other tested measures, we performed a one-way analysis of variance (ANOVA) across correlation values (separately for target power and ISPC). For the ROIs in which the ANOVA indicated a significant difference between the alignment to different mechanisms, we assessed the significance of the mean difference between the top two communication principles (*⍺* = 0.05) (Fig.2D-centre,bottom; Fig.3D-centre,bottom).

The significance of the relationship between EWC-power/ISPC correlations and functional hierarchy along the unimodal-transmodal axis was tested through spin-permutation (*S* = 10000 permutations) of the parcellated cortical maps using the technique employed in (Váša et al., 2018).

## Code accessibility

All the code used to analyse data is available at https://github.com/vmadanmohan/EWC/tree/main/flex_comm_principles.

## Acknowledgements

VMM was supported by the Melbourne Research Scholarship, University of Melbourne. AMH was supported by the Australian Research Council (DE220101019). CS acknowledges support from the Australian Research Council (DP170101815). AZ is supported by an ARC Future Fellowship (FT220100091) and the Rebecca L. Cooper Foundation. RFHC is funded by an NHMRC Emerging Leadership Investigator Grant (Grant number: 2017527). Data were provided [in part] by the Human Connectome Project, WU-Minn Consortium (Principal Investigators: David Van Essen and Kamil Ugurbil; 1U54MH091657) funded by the 16 NIH Institutes and Centers that support the NIH Blueprint for Neuroscience Research; and by the McDonnell Center for Systems Neuroscience at Washington University. This research was supported by The University of Melbourne’s Research Computing Services and the Petascale Campus Initiative.

## References

Abeysuriya, R. G., Hadida, J., Sotiropoulos, S. N., Jbabdi, S., Becker, R., Hunt, B. A. E., Brookes, M. J., & Woolrich, M. W. (2018). A biophysical model of dynamic balancing of excitation and inhibition in fast oscillatory large-scale networks. PLoS Computational Biology, 14(2), 1–27. 10.1371/journal.pcbi.1006007

Akam, T., & Kullmann, D. M. (2010). Oscillations and filtering networks support flexible routing of information. Neuron, 67(2), 308–320. 10.1016/j.neuron.2010.06.019

Avena-Koenigsberger, A., Misic, B., & Sporns, O. (2018). Communication dynamics in complex brain networks. In Nature Reviews Neuroscience (Vol. 19, Issue 1, pp. 17–33). Nature Publishing Group. 10.1038/nrn.2017.149

Avena-Koenigsberger, A., Yan, X., Kolchinsky, A., Van Den Heuvel, M. P., Hagmann, P., & Sporns, O. (2019). A spectrum of routing strategies for brain networks. PLoS Computational Biology, 15(3), 1–24. 10.1371/journal.pcbi.1006833

Başar, E., Başar-Eroglu, C., Karakaş, S., & Schürmann, M. (2001). Gamma, alpha, delta, and theta oscillations govern cognitive processes. International Journal of Psychophysiology, 39, 241–248.

Bazinet, V., Vos de Wael, R., Hagmann, P., Bernhardt, B. C., & Misic, B. (2021). Multiscale communication in cortico-cortical networks. NeuroImage, 243. 10.1016/j.neuroimage.2021.118546

Betzel, R. F., Griffa, A., Hagmann, P., & Mišić, B. (2018). Distance-dependent consensus thresholds for generating group-representative structural brain networks. Network Neuroscience, 3(2), 475–496. 10.1162/netn_a_00075

Bonnefond, M., Kastner, S., & Jensen, O. (2017). Communication between brain areas based on nested oscillations. ENeuro, 4(2), 1–14. 10.1523/ENEURO.0153-16.2017

Bosman, C. A., Schoffelen, J. M., Brunet, N., Oostenveld, R., Bastos, A. M., Womelsdorf, T., Rubehn, B., Stieglitz, T., De Weerd, P., & Fries, P. (2012). Attentional Stimulus Selection through Selective Synchronization between Monkey Visual Areas. Neuron, 75(5), 875–888. 10.1016/j.neuron.2012.06.037

Bruña, R., Maestú, F., & Pereda, E. (2018). Phase locking value revisited: Teaching new tricks to an old dog. Journal of Neural Engineering, 15(5). 10.1088/1741-2552/aacfe4

Buehlmann, A., & Deco, G. (2008). The neuronal basis of attention: Rate versus synchronization modulation. Journal of Neuroscience, 28(30), 7679–7686. 10.1523/JNEUROSCI.5640-07.2008

Buehlmann, A., & Deco, G. (2010). Optimal information transfer in the cortex through synchronization. PLoS Computational Biology, 6(9). 10.1371/journal.pcbi.1000934

Buzsáki, G., & Draguhn, A. (2004). Neuronal Oscillations in Cortical Networks. Science, 304, 1926–1929. www.sciencemag.org

Chapeton, J. I., Haque, R., Witg, J. H., Inati, S. K., & Zaghloul, K. A. (2019). Large-Scale Communication in the Human Brain Is Rhythmically Modulated through Alpha Coherence. Current Biology, 29(17), 2801–2811.e5. 10.1016/j.cub.2019.07.014

Cohen, J. R., & D’Esposito, M. (2016). The segregation and integration of distinct brain networks and their relationship to cognition. Journal of Neuroscience, 36(48), 12083–12094. 10.1523/JNEUROSCI.2965-15.2016

Cohen, M. X. (2014). Analyzing Neural Time Series Data (J. Grafman, Ed.). The MIT Press.

Colclough, G. L., Brookes, M. J., Smith, S. M., & Woolrich, M. W. (2015). A symmetric multivariate leakage correction for MEG connectomes. NeuroImage, 117, 439–448. 10.1016/j.neuroimage.2015.03.071

Dale, A. M., Liu, A. K., Fischl, B. R., Buckner, R. L., Belliveau, J. W., Lewine, J. D., & Halgren, E. (2000). Dynamic Statistical Parametric Mapping: Combining fMRI and MEG for High-Resolution Imaging of Cortical Activity. Neuron, 26, 55–67.

Deco, G., Jirs, V., McIntosh, A. R., Sporns, O., & Kötter, R. (2009). Key role of coupling, delay, and noise in resting brain fluctuations. Proceedings of the National Academy of Sciences of the United States of America, 106(25), 10302–10307. 10.1073/pnas.0901831106

Destrieux, C., Fischl, B., Dale, A., & Halgren, E. (2010). Automatic parcellation of human cortical gyri and sulci using standard anatomical nomenclature. NeuroImage, 53(1), 1–15. 10.1016/j.neuroimage.2010.06.010

Engel, A. K., & Fries, P. (2010). Beta-band oscillations-signalling the status quo? In Current Opinion in Neurobiology (Vol. 20, Issue 2, pp. 156–165). Elsevier Ltd. 10.1016/j.conb.2010.02.015

Engel, A. K., Fries, P., & Singer, W. (2001). Dynamic Predictions: Oscillations and Synchrony in Top-Down Processing. Nature Reviews Neuroscience, 2. www.nature.com/reviews/neuro

Friederici, A. D. (2011). THE BRAIN BASIS OF LANGUAGE PROCESSING: FROM STRUCTURE TO FUNCTION. Physiol Rev, 91, 1357–1392. 10.1152/physrev.00006.2011.-Lan

Fries, P. (2005). A mechanism for cognitive dynamics: Neuronal communication through neuronal coherence. Trends in Cognitive Sciences, 9(10), 474–480. 10.1016/j.Lcs.2005.08.011

Fries, P. (2009). Neuronal gamma-band synchronization as a fundamental process in cortical computation. Annual Review of Neuroscience, 32, 209–224. 10.1146/annurev.neuro.051508.135603

Fries, P. (2015). Rhythms for Cognition: Communication through Coherence. Neuron, 88(1), 220–235. 10.1016/j.neuron.2015.09.034

Girardi-Schappo, M., Fadaie, F., Lee, H. M., Caldairou, B., Sziklas, V., Crane, J., Bernhardt, B. C., Bernasconi, A., & Bernasconi, N. (2021). Altered communication dynamics reflect cognitive deficits in temporal lobe epilepsy. Epilepsia, 62(4), 1022–1033. 10.1111/epi.16864

Glasser, M. F., Sotiropoulos, S. N., Wilson, J. A., Coalson, T. S., Fischl, B., Andersson, J. L., Xu, J., Jbabdi, S., Webster, M., Polimeni, J. R., Van Essen, D. C., & Jenkinson, M. (2013). The minimal preprocessing pipelines for the Human Connectome Project. NeuroImage, 80, 105–124. 10.1016/j.neuroimage.2013.04.127

Griffa, A., Mach, M., Dedelley, J., Gutierrez-Barragan, D., Gozzi, A., Allali, G., Grandjean, J., Van De Ville, D., & Amico, E. (2023). Evidence for increased parallel information transmission in human brain networks compared to macaques and male mice. Nature Communications, 14(1), 8216. 10.1038/s41467-023-43971-z

Griffa, A., Ricaud, B., Benzi, K., Bresson, X., Daducci, A., Vandergheynst, P., Thiran, J. P., & Hagmann, P. (2017). Transient networks of spatio-temporal connectivity map communication pathways in brain functional systems. NeuroImage, 155, 490–502. 10.1016/j.neuroimage.2017.04.015

Hahn, G., Bujan, A. F., Frégnac, Y., Aertsen, A., & Kumar, A. (2014). Communication through Resonance in Spiking Neuronal Networks. PLoS Computational Biology, 10(8). 10.1371/journal.pcbi.1003811

Hahn, G., Ponce-Alvarez, A., Deco, G., Aertsen, A., & Kumar, A. (2019). Portraits of communication in neuronal networks. Nature Reviews Neuroscience, 20(2), 117–127. 10.1038/s41583-018-0094-0

Hanslmayr, S., Staudigl, T., & Fellner, M. C. (2012). Oscillatory power decreases and long-term memory: The information via desynchronization hypothesis. In Frontiers in Human Neuroscience (Issue MARCH 2012, pp. 1–20). Frontiers Media S. A. 10.3389/fnhum.2012.00074

Hickok, G., & Poeppel, D. (2007). The cortical organization of speech processing. Nature Reviews Neuroscience, 8, 393–402. www.nature.com/reviews/neuro

Hillebrand, A., Tewarie, P., Van Dellen, E., Yu, M., Carbo, E. W. S., Douw, L., Gouw, A. A., Van Straaten, E. C. W., & Stam, C. J. (2016). Direction of information flow in large-scale resting-state networks is frequency-dependent. Proceedings of the National Academy of Sciences of the United States of America, 113(14), 3867–3872. 10.1073/pnas.1515657113

Jensen, O., & Mazaheri, A. (2010). Shaping functional architecture by oscillatory alpha activity: Gating by inhibition. Frontiers in Human Neuroscience, 4. 10.3389/fnhum.2010.00186

Klimesch, W., Sauseng, P., & Hanslmayr, S. (2007). EEG alpha oscillations: The inhibition-timing hypothesis. Brain Research Reviews, 53(1), 63–88. 10.1016/j.brainresrev.2006.06.003

Lachaux, J. P., Rodriguez, E., Martinerie, J., & Varela, F. J. (1999). Measuring phase synchrony in brain signals. Human Brain Mapping, 8(4), 194–208. 10.1002/(SICI)1097-0193(1999)8:4<194::AID-HBM4>3.0.CO;2-C

Larson-Prior, L. J., Oostenveld, R., Della Penna, S., Michalareas, G., Prior, F., Babajani-Feremi, A., Schoffelen, J. M., Marzet, L., de Pasquale, F., Di Pompeo, F., Stout, J., Woolrich, M., Luo, Q., Bucholz, R., Fries, P., Pizzella, V., Romani, G. L., Corbetta, M., & Snyder, A. Z. (2013). Adding dynamics to the Human Connectome Project with MEG. NeuroImage, 80, 190–201. 10.1016/j.neuroimage.2013.05.056

Lega, B., Burke, J., Jacobs, J., & Kahana, M. J. (2016). Slow-Theta-to-Gamma Phase-Amplitude Coupling in Human Hippocampus Supports the Formation of New Episodic Memories. Cerebral Cortex, 26(1), 268–278. 10.1093/cercor/bhu232

Lerner, T. N., Ye, L., & Deisseroth, K. (2016). Communication in Neural Circuits: Tools, Opportunities, and Challenges. Cell, 164(6), 1136–1150. 10.1016/j.cell.2016.02.027

Madan Mohan, V., Varley, T. F., Cash, R. F. H., Seguin, C., & Zalesky, A. (2025). Event-Marked Windowed Communication: Inferring Activity Propagation from Neural Time Series. Human Brain Mapping, 46(8). 10.1002/hbm.70223

Mansour L, S., Tian, Y., Yeo, B. T. T., Cropley, V., & Zalesky, A. (2021). High-resolution connectomic fingerprints: Mapping neural identity and behavior. NeuroImage, 229. 10.1016/j.neuroimage.2020.117695

Margulies, D. S., Ghosh, S. S., Goulas, A., Falkiewicz, M., Huntenburg, J. M., Langs, G., Bezgin, G., Eickhoff, S. B., Castellanos, F. X., Petrides, M., Jefferies, E., & Smallwood, J. (2016). Situating the default-mode network along a principal gradient of macroscale cortical organization. Proceedings of the National Academy of Sciences of the United States of America, 113(44), 12574–12579. 10.1073/pnas.1608282113

Mesulam, M.-M. (1998). From sensation to cognition. Brain, 121, 1013–1052.

Milner, A. D., & Goodale, M. A. (1992). Separate visual pathways for perception and action. In Trends in Neurosciences (Vol. 15, pp. 20–25).

Niso, G., Rogers, C., Moreau, J. T., Chen, L. Y., Madjar, C., Das, S., Bock, E., Tadel, F., Evans, A. C., Jolicoeur, P., & Baillet, S. (2016). OMEGA: The Open MEG Archive. NeuroImage, 124, 1182–1187. 10.1016/j.neuroimage.2015.04.028

Niso, G., Tadel, F., Bock, E., Cousineau, M., Santos, A., & Baillet, S. (2019). Brainstorm pipeline analysis of resting-state data from the open MEG archive. Frontiers in Neuroscience, 13(APR). 10.3389/fnins.2019.00284

O’Connor, D. H., Fukui, M. M., Pinsk, M. A., & Kastner, S. (2002). Attention modulates responses in the human lateral geniculate nucleus. Nature Neuroscience, 5(11), 1203–1209. 10.1038/nn957

Odean, N. N., Sanayei, M., & Shadlen, M. N. (2023). Transient Oscillations of Neural Firing Rate Associated With Routing of Evidence in a Perceptual Decision. The Journal of Neuroscience, 43(37), 6369–6383. 10.1523/JNEUROSCI.2200-22.2023

O’Neill, G. C., Bauer, M., Woolrich, M. W., Morris, P. G., Barnes, G. R., & Brookes, M. J. (2015). Dynamic recruitment of resting state sub-networks. NeuroImage, 115, 85–95. 10.1016/j.neuroimage.2015.04.030

Osipova, D., Hermes, D., & Jensen, O. (2008). Gamma power is phase-locked to posterior alpha activity. PLoS ONE, 3(12). 10.1371/journal.pone.0003990

Palmigiano, A., Geisel, T., Wolf, F., & Battaglia, D. (2017). Flexible information routing by transient synchrony. Nature Neuroscience, 20(7), 1014–1022. 10.1038/nn.4569

Papadopoulos, L., Lynn, C. W., Battaglia, D., & Bassett, D. S. (2020). Relations between large-scale brain connectivity and effects of regional stimulation depend on collective dynamical state. PLoS Computational Biology, 16(9). 10.1371/journal.pcbi.1008144

Quinn, A. J., van Es, M. W. J., Gohil, C., & Woolrich, M. W. (2023). OHBA SoWware Library in Python (OSL) (0.2.0).

Reyes, A. D. (2003). Synchrony-dependent propagation of firing rate in iteratively constructed networks in vitro. Nature Neuroscience, 6(6), 593. http://www.nature.com/natureneuroscience

Roland, P. E., Hilgetag, C. C., & Deco, G. (2014). Cortico-cortical communication dynamics. In *Frontiers in Systems Neuroscience* (Vol. 8, Issue MAY). Frontiers Research Foundation. 10.3389/fnsys.2014.00019

Royer, J., Rodríguez-Cruces, R., Tavakol, S., Larivière, S., Herholz, P., Li, Q., Vos de Wael, R., Paquola, C., Benkarim, O., Park, B. yong, Lowe, A. J., Margulies, D., Smallwood, J., Bernasconi, A., Bernasconi, N., Frauscher, B., & Bernhardt, B. C. (2022). An Open MRI Dataset For Multiscale Neuroscience. Scientific Data, 9(1). 10.1038/s41597-022-01682-y

Schaefer, A., Kong, R., Gordon, E. M., Laumann, T. O., Zuo, X.-N., Holmes, A. J., Eickhoff, S. B., & Yeo, B. T. T. (2018). Local-Global Parcellation of the Human Cerebral Cortex from Intrinsic Functional Connectivity MRI. Cerebral Cortex, 28(9), 3095–3114. 10.1093/cercor/bhx179

Schipul, S. E., Keller, T. A., & Just, M. A. (2011). Inter-regional brain communication and its disturbance in autism. Frontiers in Systems Neuroscience, 5. 10.3389/fnsys.2011.00010

Schnitzler, A., & Gross, J. (2005). Normal and pathological oscillatory communication in the brain. Nature Reviews Neuroscience, 6(4), 285–296. 10.1038/nrn1650

Seguin, C., Jedynak, M., David, O., Mansour, S., Sporns, O., & Zalesky, A. (2023). Communication dynamics in the human connectome shape the cortex-wide propagation of direct electrical stimulation. Neuron. 10.1016/j.neuron.2023.01.027

Seguin, C., Mansour L, S., Sporns, O., Zalesky, A., & Calamante, F. (2022). Network communication models narrow the gap between the modular organization of structural and functional brain networks. NeuroImage, 257. 10.1016/j.neuroimage.2022.119323

Seguin, C., Sporns, O., & Zalesky, A. (2023). Brain network communication: concepts, models and applications. Nature Reviews Neuroscience. 10.1038/s41583-023-00718-5

Seguin, C., Tian, Y., & Zalesky, A. (2020). Network communication models improve the behavioral and functional predictive utility of the human structural connectome. Network Neuroscience, 4(4), 980–1006. 10.1162/netn_a_00161

Smith, R. E., Tournier, J. D., Calamante, F., & Connelly, A. (2012). Anatomically-constrained tractography: Improved diffusion MRI streamlines tractography through effective use of anatomical information. NeuroImage, 62(3), 1924–1938. 10.1016/j.neuroimage.2012.06.005

Smith, S. M., Beckmann, C. F., Andersson, J., Auerbach, E. J., Bijsterbosch, J., Douaud, G., Duff, E., Feinberg, D. A., Griffanti, L., Harms, M. P., Kelly, M., Laumann, T., Miller, K. L., Moeller, S., Petersen, S., Power, J., Salimi-Khorshidi, G., Snyder, A. Z., Vu, A. T., … Glasser, M. F. (2013). Resting-state fMRI in the Human Connectome Project. NeuroImage, 80, 144–168. 10.1016/j.neuroimage.2013.05.039

Sorrentino, P., Seguin, C., Rucco, R., LiparoL, M., Lopez, E. T., Bonavita, S., Quarantelli, M., Sorrentino, G., Jirsa, V., & Zalesky, A. (2021). The structural connectome constrains fast brain dynamics. ELife, 10. 10.7554/etiife.67400

Tadel, F., Baillet, S., Mosher, J. C., Pantazis, D., & Leahy, R. M. (2011). Brainstorm: A user-friendly application for MEG/EEG analysis. Computational Intelligence and Neuroscience, 2011. 10.1155/2011/879716

The MathWorks Inc. (2022). *MATLAB version: 9.13.0 (R2022b)* (R2022b). The MathWorks Inc.

Tournier, J. D., Calamante, F., & Connelly, A. (2007). Robust determination of the fibre orientation distribution in diffusion MRI: Non-negativity constrained super-resolved spherical deconvolution. NeuroImage, 35(4), 1459–1472. 10.1016/j.neuroimage.2007.02.016

Tournier, J. D., Calamante, F., & Connelly, A. (2010). Improved probabilistic streamlines tractography by 2nd order integration over fibre orientation distributions. Proceedings of the International Society for Magnetic Resonance in Medicine, 1670.

Tournier, J. D., Calamante, F., & Connelly, A. (2012). MRtrix: Diffusion tractography in crossing fiber regions. International Journal of Imaging Systems and Technology, 22(1), 53–66. 10.1002/ima.22005

Uusitalo, M. A., & Ilmoniemi, R. J. (1997). Communication Signal-space projection method for separating MEG or EEG into components. Biol. Eng. & Comput, 35, 135–140.

Van, C., Abbott, L. F., & Ermentrout, G. B. (1994). When Inhibition not Excitation Synchronizes Neural Firing. In Journal of Computational Neuroscience (Vol. 1).

Van Der Maaten, L., Postma, E., & Van Den Herik, J. (2009). Dimensionality Reduction: A Comparative Review. http://www.uvt.nl/Lcc

Van Essen, D. C., Smith, S. M., Barch, D. M., Behrens, T. E. J., Yacoub, E., & Ugurbil, K. (2013). The WU-Minn Human Connectome Project: An overview. NeuroImage, 80, 62–79. 10.1016/j.neuroimage.2013.05.041

Váša, F., Seidlitz, J., Romero-Garcia, R., Whitaker, K. J., Rosenthal, G., Vértes, P. E., Shinn, M., Alexander-Bloch, A., Fonagy, P., Dolan, R. J., Jones, P. B., Goodyer, I. M., Sporns, O., & Bullmore, E. T. (2018). Adolescent tuning of association cortex in human structural brain networks. Cerebral Cortex, 28(1), 281–294. 10.1093/cercor/bhx249

Vázquez-Rodríguez, B., Liu, Z. Q., Hagmann, P., & Misic, B. (2020). Signal propagation via cortical hierarchies. Network Neuroscience, 4(4), 1072–1090. 10.1162/netn_a_00153

Voloh, B., & Womelsdorf, T. (2016). A role of phase-resetng in coordinating large scale neural networks during attention and goal-directed behavior. Frontiers in Systems Neuroscience, 10(MAR). 10.3389/fnsys.2016.00018

Von Stein, A., & Sarnthein, J. (2000). Different frequencies for different scales of cortical integration: from local gamma to long range alpha/theta synchronization. International Journal of Psychophysiology, 38, 301313.

Womelsdorf, T., Schoffelen, J.-M., Oostenveld, R., Singer, W., Desimone, R., Engel, A. K., & Fries, P. (2007). Modulation of Neuronal Interactions through Neuronal Synchronization. New Series, 316(5831), 1609–1612. 10.1126/science.ll39178

Zalesky, A., Fornito, A., Cocchi, L., Gollo, L. L., van den Heuvel, M. P., & Breakspear, M. (2016). Connectome sensitivity or specificity: which is more important? NeuroImage, 142, 407–420. 10.1016/j.neuroimage.2016.06.035

