## Supplementary Figures for "Evaluating oscillatory mechanisms underlying flexible neural communication in the human brain"

# S1.

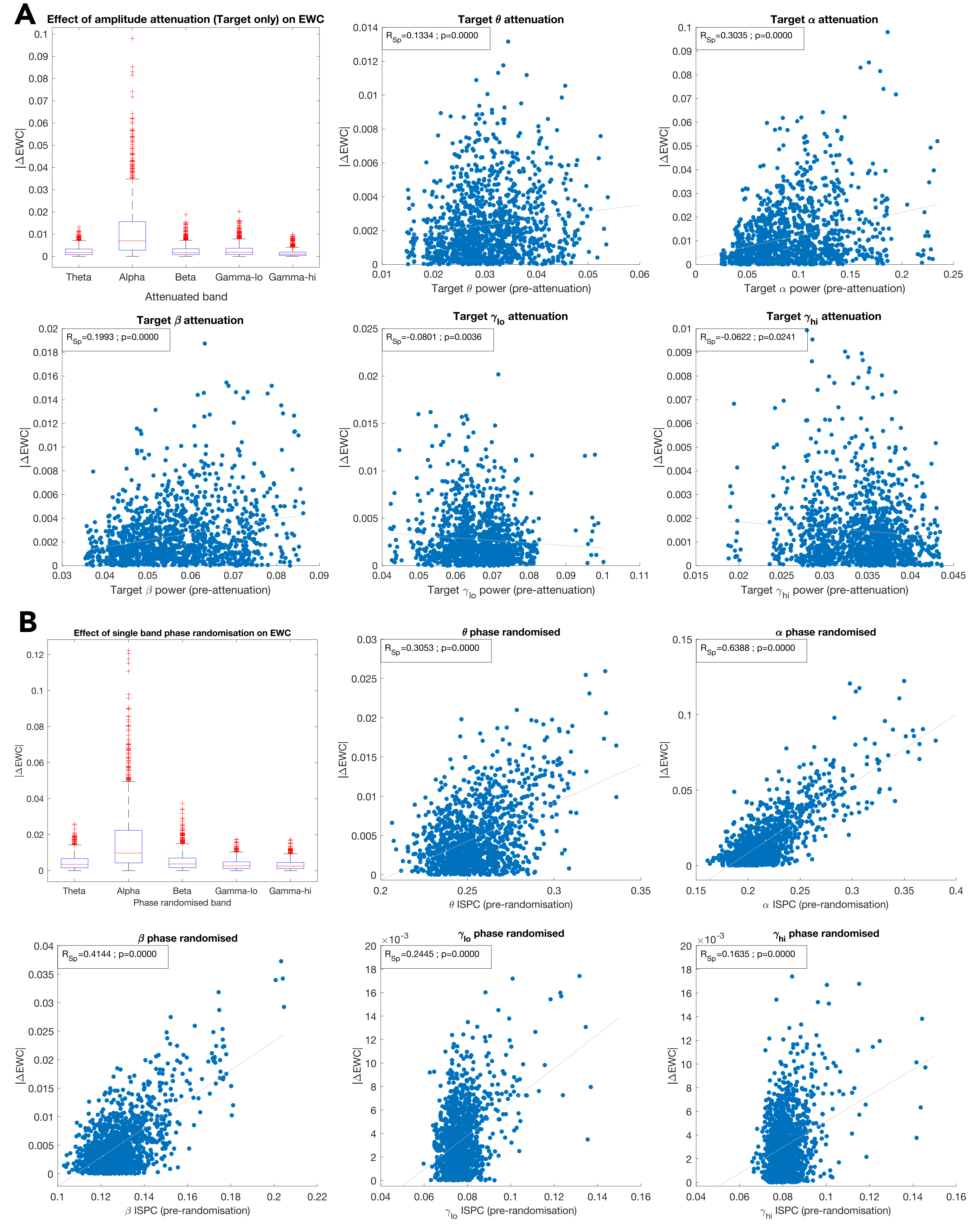


*Figure S1 - Trivial dependence of EWC (partial correlation) on target power and intersite phase clustering (ISPC) for one subject.* ***(A)*** *We quantify EWC's trivial dependence on target power by artifically attenuating the Fourier amplitudes of each frequency band by 90%, resynthesising the amplitude-attenuated version of the signal, and computing the EWC in the modified signal. The absolute change in EWC (for each source target pair - overall distribution in boxplots) is plotted against the original power in the band prior to band-attenuation.* ***(B)*** *To quantify the dependence of EWC on the ISPC or phase locking value in the different bands, we randomised the phase relationship in each band by randomly shifting the instantaneous phases of regions relative to each other to destroy the original ISPC between them. The modified bands are reconstituted into the original signal, and the EWC is then computed on the phase modified signal. Similar to the power, the absolute change to EWC is plotted against the original value of ISPC. In both (A) and (B), the relationship between the EWC change and power/ISPC is quantified as the Spearman correlation between the variables. This relationship between the measures can be ascribed to their trivial dependence, since our artifical alteration of the oscillatory profile of the signal caused the EWC to change.*

*Crucially, being the trivial or default relationship between EWC and the neural oscillatory measures, a similar degree of association would be observed between EWC and power/ISPC when simultaneously estimated from the cyclically permuted surrogate data. Thus, any empirical relationship that significantly exceeds the cyclic-surrogate-null distribution will represent a communication principle that goes above-and-beyond the default statistical relationship between the measures.*

# S2.

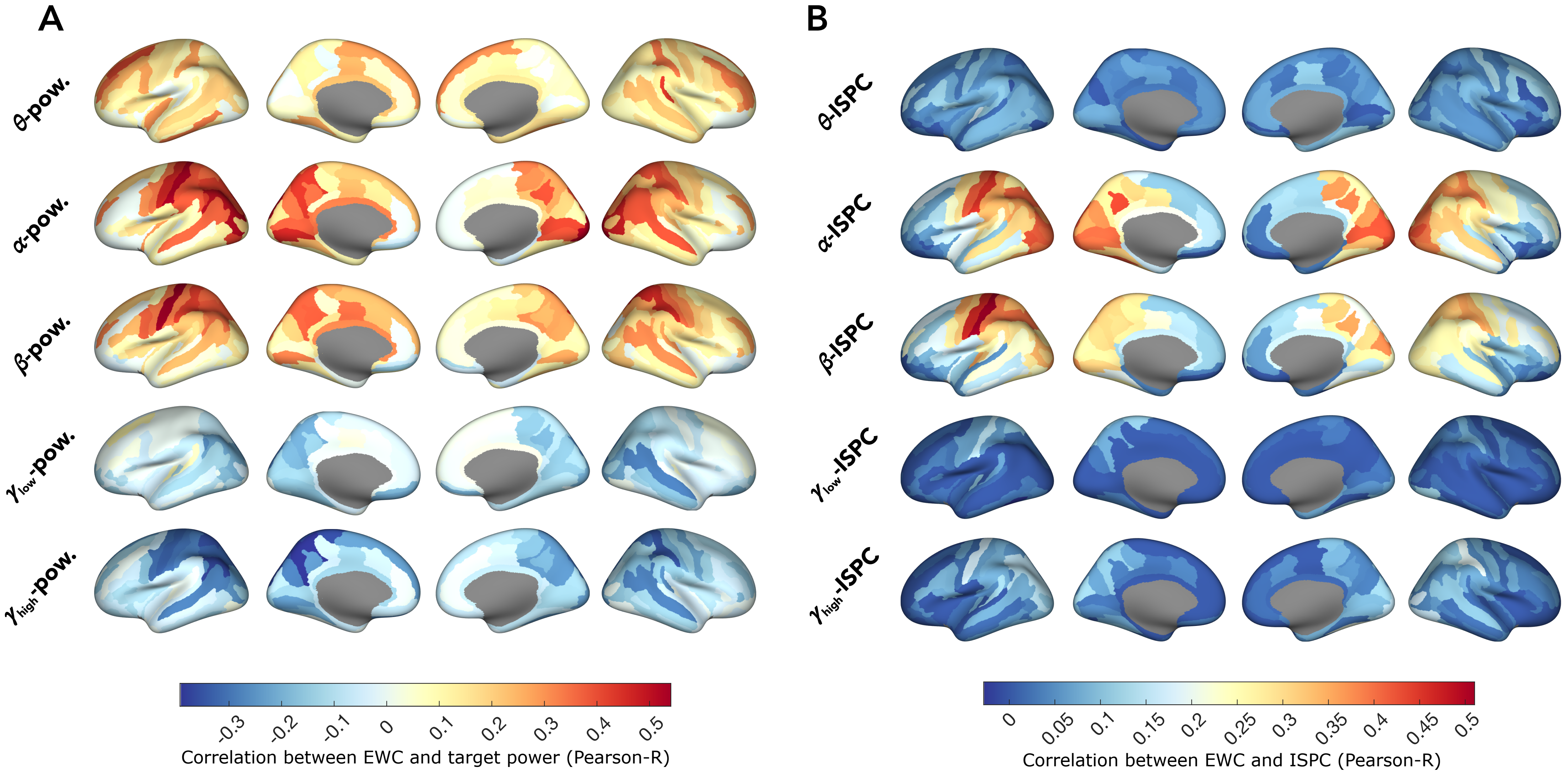


*Figure S2 - Dependence of inferred communication on* ***(A)*** *Target power; and* ***(B)*** *Intersite phase clustering (ISPC), in 5 participants from the OMEGA dataset. (A) and (B) are generated in a manner similar to Fig.2B and Fig.3B.*

S3.

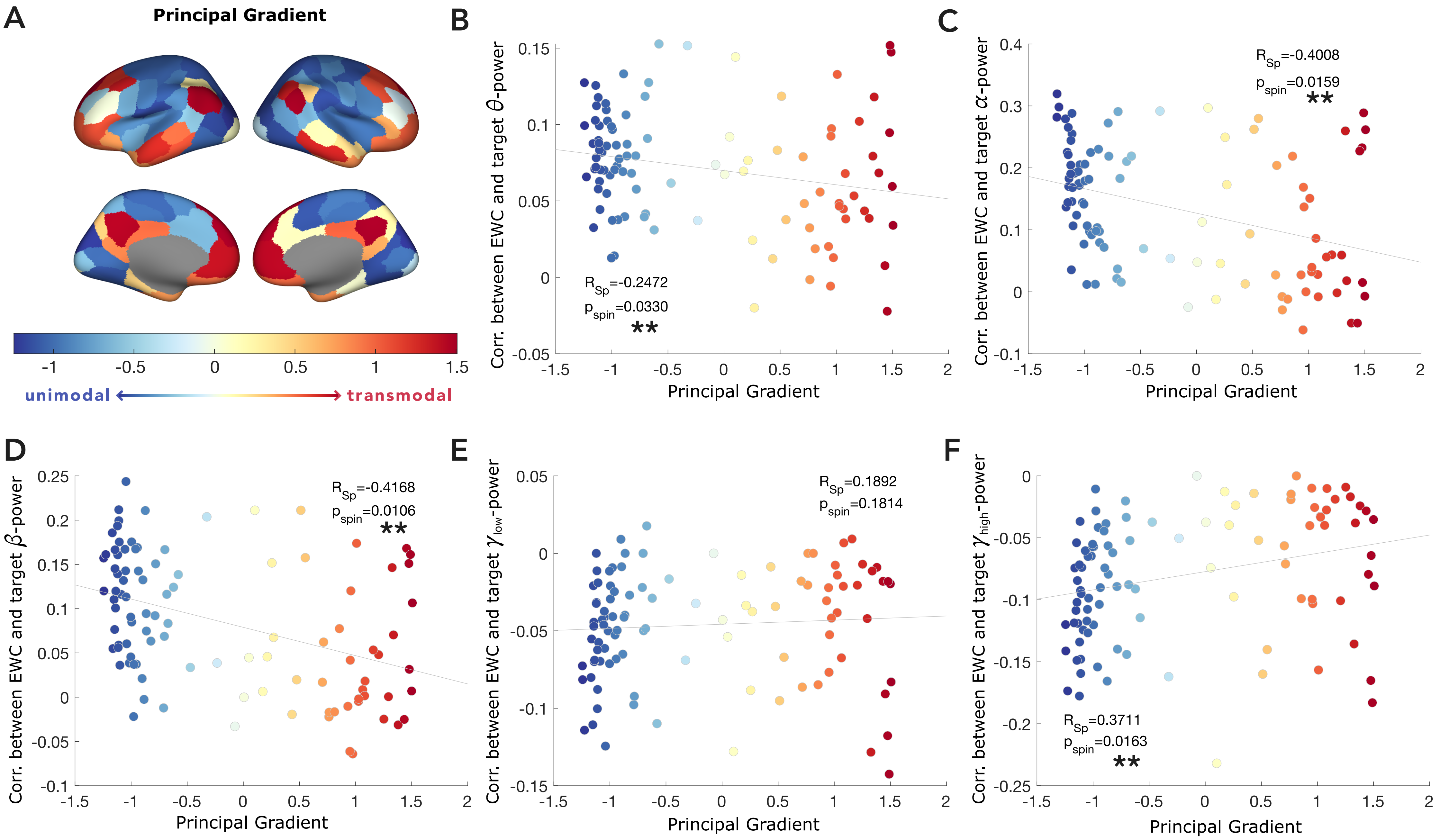


*Figure S3 - Strength of the correlation between a EWC and target power depends on the region's placement along the principal functional gradient.* **(A)** *Principal functional gradient derived from diffusion embedding of group-averaged FC.* **(B-F)** *Correlation between EWC and target power in the frequency bands of interest, as a function of the region's principal gradient coefficient (also represented in the colour of the point). Each point represents a brain region. Relationship to functional gradient is quantified as the Spearman correlation, and p-value is obtained by comparison to surrogate dataset with 10000 spin permutations.*
